# Fauxcurrence: simulating multi-species occurrences for null models in species distribution modelling and biogeography

**DOI:** 10.1101/2021.04.22.440999

**Authors:** Owen G. Osborne, Henry G. Fell, Hannah Atkins, Jan van Tol, Daniel Phillips, Leonel Herrera-Alsina, Poppy Mynard, Greta Bocedi, Cécile Gubry-Rangin, Lesley T. Lancaster, Simon Creer, Meis Nangoy, Fahri Fahri, Pungki Lupiyaningdyah, I Made Sudiana, Berry Juliandi, Justin M.J. Travis, Alexander S.T. Papadopulos, Adam C. Algar

## Abstract

Defining appropriate null expectations for species distribution hypotheses is important because sampling bias and spatial autocorrelation can produce realistic, but ecologically meaningless, geographic patterns. Generating null species occurrences with similar spatial structure to observed data can help overcome these problems, but existing methods focus on single or pairs of species and do not incorporate between-species spatial structure that may occlude comparative biogeographic analyses. Here, we describe an algorithm for generating randomised species occurrence points that mimic the within- and between-species spatial structure of real datasets and implement it in a new R package - *fauxcurrence*. The algorithm can be implemented on any geographic domain for any number of species, limited only by computing power. To demonstrate its utility, we apply the algorithm to two common analysis-types: testing the fit of species distribution models (SDMs) and evaluating niche-overlap. The method works well on all tested datasets within reasonable timescales. We found that many SDMs, despite a good fit to the data, were not significantly better than null expectations and identified only two cases (out of a possible 32) of significantly higher niche divergence than expected by chance. The package is user-friendly, flexible and has many potential applications beyond those tested here, such as joint SDM evaluation and species co-occurrence analysis, spanning the areas of ecology, evolutionary biology and biogeography.

## 1 INTRODUCTION

Eco-geographical hypothesis testing using species occurrence data can be hampered by spatial autocorrelation and the difficulty of defining appropriate null expectations (Bahn & Mcgill, 2007; Beale, Lennon, & Gimona, 2008; Chapman, 2010; Fourcade, Besnard, & Secondi, 2018; Moore, Bagchi, Aiello-Lammens, & Schlichting, 2018). A major issue is that spatial clustering of conspecific, or separation of heterospecific, occurrence records can be affected by multiple factors, which are often difficult to disentangle. These include: i) habitat suitability (Phillips, Anderson, & Schapire, 2006), ii) dispersal limitation (Glor & Warren, 2010), iii) interactions between individuals of the same or different species such as conspecific attraction, competitive exclusion or mutualism (Mielke et al., 2020), or iv) sampling bias, where occurrence records are more likely to be collected from more easily accessible or intensively studied areas (Phillips et al., 2009).

One approach to overcome these issues has been to use null species occurrences to define the expectations if only the inherent spatial structure within species has shaped their distributions (Beale et al., 2008; Algar, Mahler, Glor, & Losos, 2013). To define the null expectation, these approaches use an iterative procedure to produce null species distributions with similar spatial structure to observed occurrences, excluding any consideration of environment or specific geographic location. While these methods are well-suited to testing habitat-suitability hypotheses for single species, they do not take spatial structure between species into account, making them unsuitable for testing multispecies hypotheses involving, for example, niche overlap or range boundaries. Some methods designed to test niche overlap hypotheses employ null models for pairs of species, but these either do not take spatial structure into account (Warren, Glor, & Turelli, 2008), or simply translocate the entire set of occurrence points. This preserves spatial structure but limits their application to species which are range-restricted relative to the study region (Nunes & Pearson, 2017).

Here, we present a method to fill this gap, which is implemented in a new *R* package – *fauxcurrence* version 1.0 (available at https://github.com/ogosborne/fauxcurrence). The package can produce null species occurrences which preserve the spatial structure within and between an arbitrary number of species, and provides many options to tailor these occurrences to the user’s needs. We demonstrate the utility of the package using a dataset of 22 species of plants, vertebrates and arthropods. We use the resulting null occurrence points to test the significance of species distribution models (SDMs) and to test for significant deviations from null expectations of niche overlap between species.

## 2 MATERIALS AND METHODS

### 2.1 Method description

Our method (Fig. 1a) has three main modes of operation, distinguished by how inter-point distances are used to define spatial structure. Within-species distances (divided into one subset per species) are always included and can also be used alone (which we refer to here as the “*Intra*” null model; Fig. 1b). The total set of between-species distances can be divided into subsets in two ways: either as sets of general between-species distances per species (i.e., the distances from a species’ occurrence points to all heterospecific occurrence points; the “*Inter*” null model; Fig. 1b) or as a separate set of distances between each pair of species in the dataset (the “*Inter-sep*” model; Fig. 1b; Appendix 1.1).

**Figure 1.**
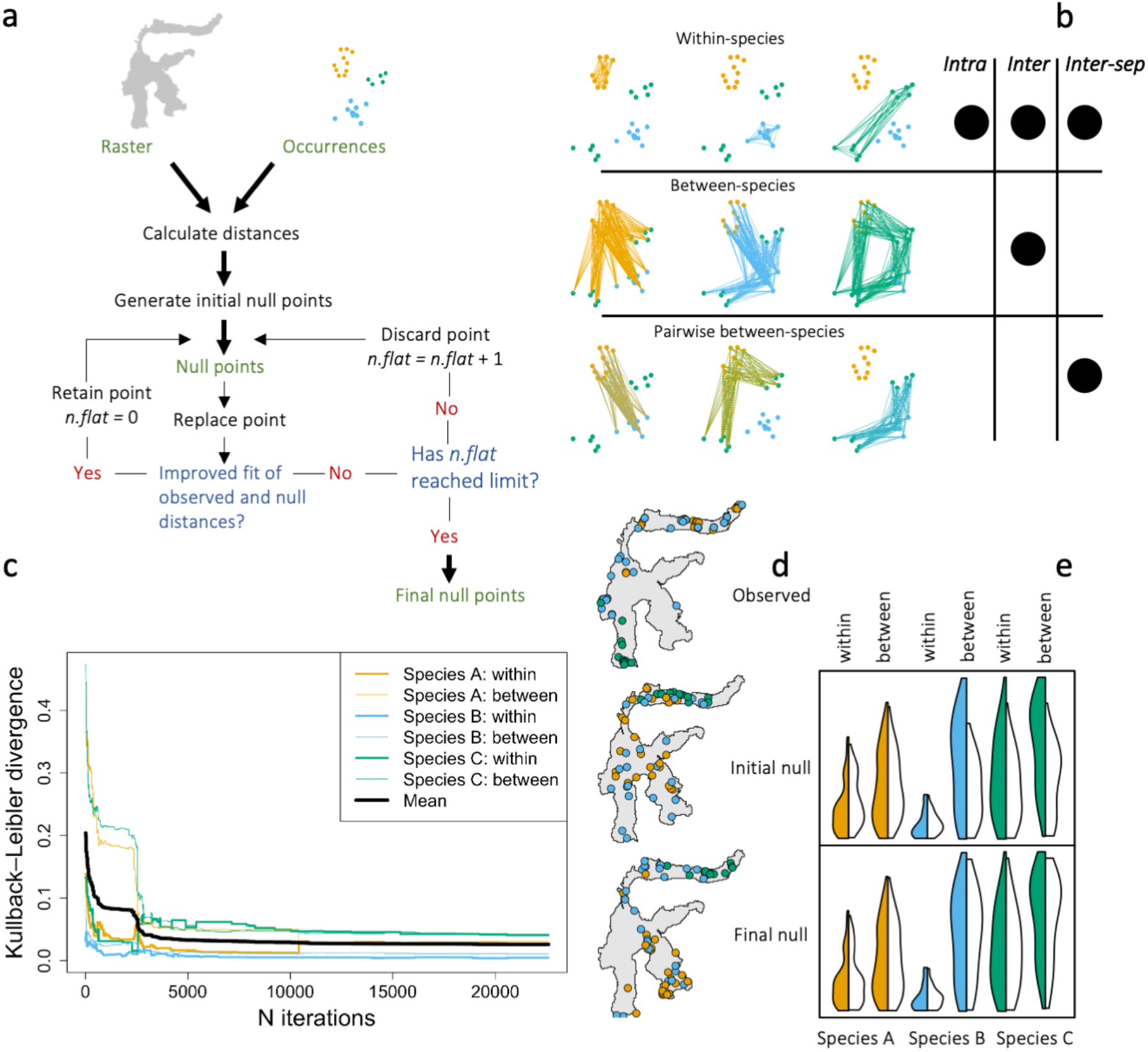
Overview of the method. A flowchart (a) shows the basic functioning of the algorithm: initial null points are generated based on observed inter-point distances, these are then iteratively improved. The algorithm finishes when the number of iterations with no improvement (*n*.*flat*) reaches a user-defined limit. There are three classes of inter-point distance sets in the algorithm (b), within-species (used in all models), general between-species (used in the *Inter* model) and pairwise between-species (used in the *Inter-sep* model). The circles to the right of each indicate which null models they are used in. Panels (c-e) show an example run of the *Inter* model. Kullback-Leibler divergence decreases across iterations (c) and this improvement can be clearly seen when comparing the initial and final null occurrence points (d; map panels). The left of each split violin plot (e) shows the density of observed interpoint distances for each of the distance sets in the model and the right shows those of the null, which are more similar to the observed distances in the final null (bottom) than the initial null (top).

The user provides a set of species occurrence points and a raster defining the study area. The algorithm begins by randomly generating one simulated occurrence point per species. It then adds occurrence points for each species by drawing each new point *D* distance away from a random existing conspecific point (where *D* is sampled from the empirical distribution function of observed within-species distances) until each species has the same number of occurrences as in the observed dataset (Appendix 1.7).

Once the initial set of simulated points are generated, the fit of their spatial structure to that of the observed points is iteratively improved. For each iteration, one point is replaced and the match between null and observed interpoint distances is evaluated using discrete Kullback-Leibler (KL) divergence (Kullback & Leibler, 1951), where smaller values indicate a better match between the simulated and observed points. Since there are multiple interpoint distance distributions (i.e., within- and between-species distances for multiple species or pairs of species), a weighted mean of KL-divergence across all distributions is used, weighted such that within- and between-species distances contribute equally. The point replacement is only retained if it improves the match, and this procedure is repeated until either no improvement has been made for a set number of iterations or a maximum iteration limit has reached (Fig. 1c-e; see Appendix 1 for full details).

### 2.2 Test data

We tested the method on seven species occurrence datasets from Sulawesi, Indonesia each containing between one and six species from a single genus (Fig. S1a; Appendix 2). We ran the *Intra* model on all datasets, and the *Inter* model on all datasets with over one species. Since the *Inter* and *Inter-sep* models are identical for species pairs, we only ran the *Inter-sep* model for datasets with over two species. The iterative improvement was continued until there had been no improvement for 10,000 iterations. For each dataset/model combination, we produced 1,000 independent, null occurrence replicates, each with similar spatial structure, but differing in the final locations selected by the algorithm.

### 2.3 SDM model-fitting and niche overlap

We used *Maxent* v. 3.4.1 (Phillips et al., 2006) to build Species Distribution Models (SDMs) for all species using the 19 *BIOCLIM* climate variables and altitude from the *WorldClim1* database (Hijmans, Cameron, Parra, Jones, & Jarvis, 2005). To determine if SDMs for the observed data had a significantly better fit than the null models, we calculated P-values by comparison of area under the receiver operating characteristic curve (AUC) in SDMs built from observed data to those from all null model replicates. For datasets with more than one species, we compared null and observed niche overlap using Schoener’s *D* (Schoener, 1968) and Warren’s *I* (Warren et al., 2008), between all pairs of congeneric species (See Appendix 3 for full details).

## 3 RESULTS

### 3.1 Method performance

Average time per iteration ranged from 3.2 milliseconds (ms) to 21.5 ms and the mean number of iterations ranged from 48,155 to 341,610 (Fig. S1). Number of occurrences was the best predictor of number of iterations to model completion, although number of species also had an effect (Table S1). Plotting KL divergence across iterations suggested that 10,000 iterations without improvement was more than sufficient to minimise the KL-divergence statistic for most datasets (Fig. S2-S4).

### 3.2 Application to SDM model-fitting and niche overlap

The AUC values were over 0.8 for 13 of 22 species and over 0.9 for four species, values typically interpreted as “excellent” or “outstanding” discrimination, respectively (Hosmer, Lemeshow, & Sturdivant, 2013). Nevertheless, just six of these were significantly greater than null expectations according to at least one of our null model types (Fig. 2a-c) and where they were significantly greater than those of one null model type, they were often not significantly different to those of the others (Fig. 2a-d). In fact, SDMs for only two species had a significantly better AUC than all applied null models (Fig. 2d).

**Figure 2.**
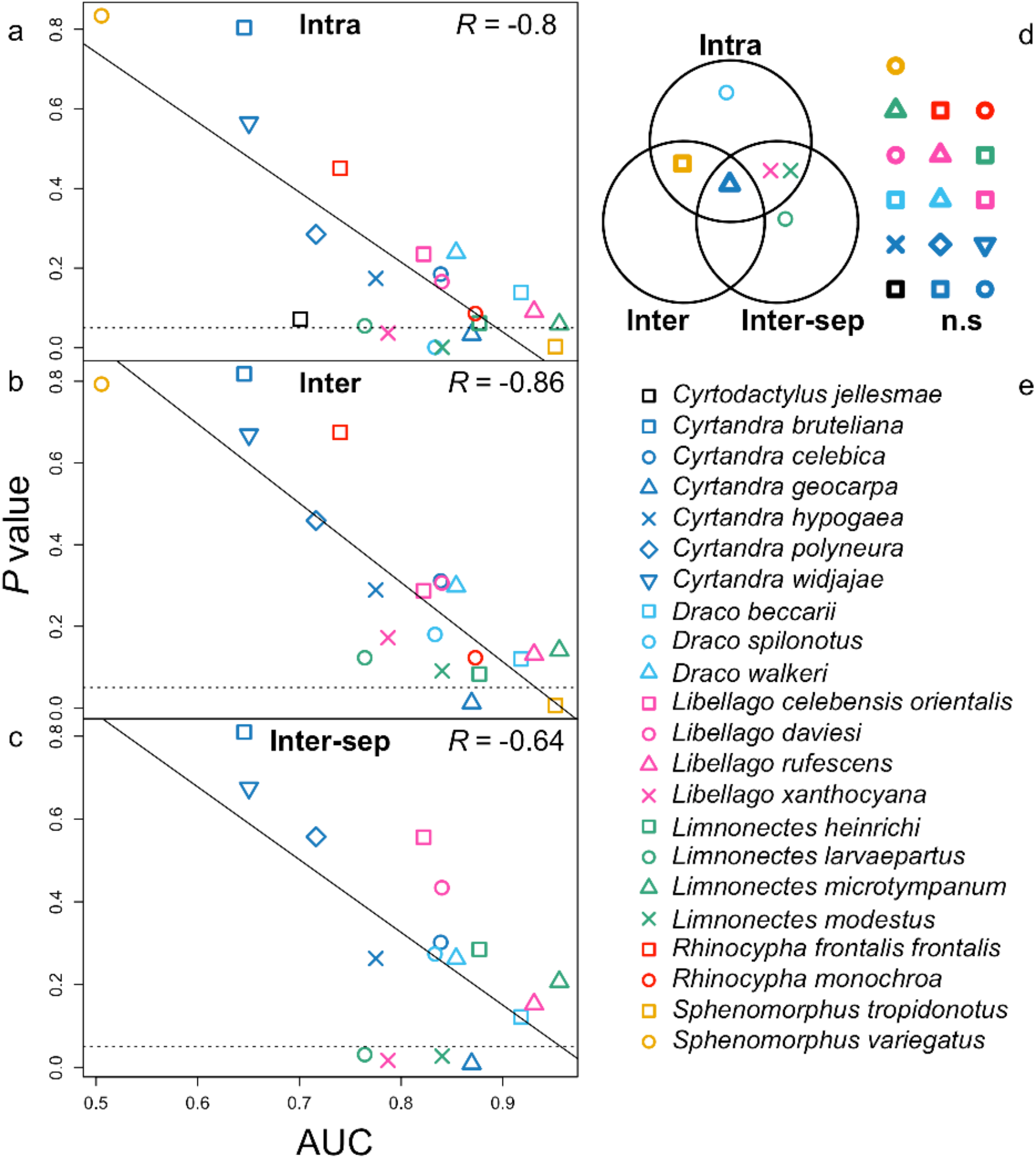
The relationship between Area Under Curve (AUC) of the Species Distribution Models (SDMs) for observed occurrences, and the *P*-values for comparison to AUC of each null model type (a-c). Dotted lines mark 0.05 on the *P-*value axis, the solid line is a linear regression line and Pearson’s correlation is shown in the top-right of each plot. A Venn diagram (d) shows the overlap in significance (*P* < 0.05) between the three models. Each labelled circle contains species with significantly higher observed AUC than those of simulated data of each null model type and those which were significantly higher than simulated data from multiple null models are shown in the appropriate intersection (those outside the circles were not significant: “n.s”). Where all implemented models agree, symbols are in bold. Species symbols are shown in the legend (e).

Observed niche divergence differed from null expectations in only two species pairs, both of which showed significant niche divergence (Fig. S5-S8). Both of these involved *Cyrtandra geocarpa*, which was also one of only two species whose SDM fit significantly better than those from all the null models (Fig. 2). Both *Inter* and *Inter-sep* null models, and both niche overlap statistics found the same two species pairs to be significant (Figs. S5-S8).

## 4 DISCUSSION

Here, we describe a new tool to help overcome the difficulties of defining null expectations when working with multi-species occurrence data. Our case study demonstrates the utility of the method. Although SDMs were calculated individually for each species, the between-species distances used in the null model had a major effect on their significance. For example, *Draco spilonotus* had a significantly better fit than the *Intra* null model (*P* = 0.001), but was not significantly different from either the *Inter* or *Inter-sep* models (*P* = 0.18 and *P* = 0.27, respectively). Such a pattern may be expected if, for example, competitive exclusion between species, rather than climate suitability, was responsible for the geographical separation of species into ranges that happen to have distinct climates (Godsoe, Franklin, & Blanchet, 2017). While AUC and *P*-values were highly correlated, many species with very high AUC scores did not have a significantly better fit than the null models. The lowest AUC score which was significantly higher than any of the null models was 0.76, underlining that SDMs with low AUC scores (e.g., < 0.75) should be treated with great caution. Using our method to demonstrate that SDMs fit significantly better than null expectations will provide much greater certainty than simple inspection of AUC.

We also demonstrated the applicability of our method to identify significant niche divergence (or niche conservatism). Our method has advantages over existing approaches. The approach of Nunes & Pearson (2017) creates null replicates by translocating and rotating observed species occurrences. This would clearly be inappropriate for a study region such as Sulawesi, since most rotations and translocations would result in a large proportion of occurrences being translocated to the sea, leading to a high number of similar replicates, as noted by the authors (Nunes & Pearson, 2017). The contorted geography of Sulawesi is not unique, and many intensely studied locations such as Isabela Island in the Galápagos archipelago and Lord Howe Island, Australia, also fall into this category. It is possible our model could have similar issues in extreme cases, where dense occurrence points cover a very large proportion of the study region. However, we expect it to work on a wider range of datasets due to our use of “as similar as possible” rather than identical spatial structure and the capability to include overland distances.

Aside from the two applications shown here, there are many other potential uses for the package. For example, the performance of joint species distribution models (Pollock et al., 2014), which jointly model environmental and community effects on species distributions, could be assessed with our approach in a similar way to our tests of SDM fit. Other potential applications include identifying significant effects of biotic factors (other species) on a focal species’ distribution (Algar et al., 2013; Giannini, Chapman, Saraiva, Alves-dos-Santos, & Biesmeijer, 2013) where the comparison of different null models can give insight into the relevance of pairwise and complex biotic interactions, and testing for significant co-occurrence of range-boundaries across clades (Swenson & Howard, 2005). More generally, *fauxcurrence*-generated occurrences could be used in any theoretical biogeographical application where realistic occurrences of species and clades are required. Overall, the method is easy to use, flexible, and can add rigour and insight into investigations of a wide range of problems in ecology, evolution and biogeography.

## ACKNOWLEDGEMENTS

The work was funded by Newton Fund (UK)/NERC (UK)/RISTEKDIKTI (Indonesia) grants awarded to JT, BJ, ACA, ASTP, CG-R, GB and LTL (Grant numbers: NE/S006923/1; NE/S006893/1; 2488/IT3.L1/PN/2020; and 3982/IT3.L1/PN/2020). GB and CG-R are funded by Royal Society University Research Fellowships (UF160614 and UF150571 respectively).

## AUTHOR CONTRIBUTIONS

Ideas for the study were conceived by ACA, OGO, HGF and ASTP. OGO wrote the package based partly on earlier code by ACA and HGF. HA, JvT and DP collated the species occurrence data. HGF and OGO ran the species distribution modelling and niche overlap analyses. ASTP and ACA supervised the project. OGO drafted the first version of the manuscript and all authors contributed to and approved the final version of the manuscript.

