## Supplementary materials for "Fauxcurrence: simulating multi-species occurrences for null models in species distribution modelling and biogeography"

**Appendix S1: additional information on options and flexibility of *fauxcurrence***

There are several available options to enhance the flexibility of the approach, detailed below.

**1.1 Distance set classes**

Whether, and how, to use between-species distances is controlled by two options: *inter.spp* and *sep.inter.spp*. If *inter.spp* is set to *FALSE*, only within-species distances are used (the *intra* model). If *inter.spp* is set to *TRUE*, the class of between-species distances used is determined by *sep.inter.spp*. If *sep.inter.spp* is *TRUE* it separates interspecific distances into all pairwise sets of species (the *Inter-sep* model. For four species, A, B, C and D, there would be six distributions of between species distances: A – B, A – C, A – D, B – C, B – D and C – D). If *sep.inter.spp* is *FALSE*, a general interspecies distance distribution for each species is used, which contains the distances from a species' occurrence points to all heterospecific occurrence points (the *Inter* null model; For species A, B, C and D, there are four distributions of between species distances in this model: A – B/C/D, B – A/C/D, C – A/B/D and D – A/B/C). While the same individual distances (i.e., the distances between every pair of interspecific occurrences) are used in both the *Inter* and *Inter-sep* models, they are divided into sets differently in the two models, and since spatial similarity between simulated and observed data is calculated by the distribution of distances in each set (see below), the two models lead to different outcomes. Thus, occurrences for each species can be simulated independently (the *Intra* model; *inter.spp=FALSE*), general proximity to heterospecifics can be taken into account (the *Inter* model; *inter.spp=TRUE*, *sep.inter.spp=FALSE*), or specific distance relationships between pairs of species can be preserved (the *Inter-sep* model; *inter.spp=TRUE*, *sep.inter.spp=TRUE*).

**1.2 Distance computation**

Distances between points can either be computed for each iteration (if *use.distmat* is set to *FALSE*) or a precomputed matrix of distances between all points in the raster can be used (if *use.distmat* is set to *TRUE*). A suitable matrix can be produced using the supplied helper function *make.distmat*, which can then be supplied to *fauxcurrence* using the *distmat* option. If *use.distmat* is *TRUE*, but *distmat* is not provided, *fauxcurrence* first calls *make.distmat* to make one. Using a precomputed distance matrix is usually faster, but the RAM requirements for larger rasters can make it computationally inviable.

**1.3 Distance measures**

Several distance measures are available, including all those from the *geosphere* (Hijmans, Williams, & Vennes, 2015) and *distRcpp* (Skinner, 2016) packages, overland distance and cost distance (implemented via the *gdistance* package (van Etten, 2017)). In fact, any function which takes a two-column matrix of longitude and latitude and outputs a vector of distances can be used, or alternatively a matrix of distances between all cells in the raster can be provided by the user (see above). The distance measure used is controlled by two options: *dist.meth* and *dist.fun*. The option *dist.meth* indicates the package used to compute distances and can be either *"distRcpp"* (for the *dist_mtom* function in the *distRcpp* package (Skinner, 2016)), *"distm"* (for the *distm* function in the *geosphere* package (Hijmans et al., 2015)) or *"costdist"* (for the cost distance method using the *geoCorrection* and *costDistance* functions in the *gdistance* package (van Etten, 2017)). The *"costdist"* method can only be used when *use.distmat* is set to *TRUE*. The option *dist.fun* sets the distance calculation to be used within the *"distm"* and *"distRcpp"* methods. For *"distRcpp"*, this should either be *"Haversine"* or *"Vincenty"*. For *"distm"*, it can be the name of any loaded function which takes the same input and produces the same output as the *distHaversine* function in *geosphere*. To use relative overland distance, as we did here, the user can choose the *"costdist"* method and supply a raster where the value of all land cells is set to the same positive value, and the value of all sea cells is set to *NA*.

**1.4 Divergence minimisation**

The KL divergence should be minimised as much as possible to produce the best possible fit between the observed and null spatial structures. Divergence tends to roughly follow an exponential decay curve as the algorithm proceeds, such that point replacements which cause a large improvement are fixed early and the curve flattens when fewer, and only minor, improvements are available (Fig. 1c, main text). Thus, the user should ensure that the curve has flattened. To do this, the divergence statistic should be plotted against number of iterations. A basic ASCII plot showing number of iterations versus mean KL divergence is printed to the console (or to a log file if *logfile* is set) if *verbose* is set to *TRUE*. In practice, the number of iterations before algorithm completion is controlled by two options: *iter.max.stg3* and *div.n.flat*. The number of iterations for which divergence values should remain unchanged before completion is set by *div.n.flat* and *iter.max.stg3* sets an upper limit on the number of iterations, regardless of KL divergence (it can also be set to *Inf,* i.e. infinity, to remove this limit). The number of iterations required varies widely between datasets, so we recommend plotting KL divergence curves for some test runs before setting *div.n.flat* and *iter.max.stg3* accordingly.

**1.5 Fixing initial points**

Instead of generating random initial points, the user can define the first point for each species by supplying a data frame (in the same format as the input occurrence points) with the *fix.seed.pts* option. If the option is used, these points are not replaced in later stages of the algorithm, so they can be used to define the (rough) centroids of the species distributions.

#### 1.6 Identical conspecific points

The algorithm can be used to simulate either presence-only data (where only one occurrence point for each species can occupy a single raster cell), or raw occurrence data (where there can be multiple identical conspecific occurrences). If the *allow.ident.conspec* option is set to *TRUE*, identical conspecific points are allowed, or if *FALSE* they are not. This should match the input data.

#### 1.7 Tuning and advanced options

There are a number of additional options to finetune the algorithm which can improve its speed or efficiency, listed below.

##### 1.7.1 Generating the first point for each species:

At the start of the algorithm a single null occurrence point is generated for each species. If *inter.spp* is *TRUE*, the algorithm ensures that the distances between species are within the observed range of distances (or within the central *init.range* quantile of the real distribution if *trim.init.range* is *TRUE*). If interspecific distances are outside the real distribution (or *init.range*), it iteratively improves the initial points by randomly replacing a point and rechecking the distances until they are within the desired range (n.b if *fix.seed.pts* is provided, initial point generation is skipped and these points are used instead). To prevent the algorithm becoming stuck with a set of points which make step-wise iterative improvement impossible, *iter.max.stg1* sets an upper limit on the number of iterations without improvement. If this is reached, initial point generation is restarted.

*iter.max.stg1*

The maximum number of iterations with no improvement allowed when generating the first point per species.

*trim.init.range*

Indicates whether the initial point for each species should fit within the central *init.range* quantile of the observed distribution if between-species distances rather than just the total range. This can speed up stage 2, especially if many species are present.

*init.range*

A value between 0 and 1 defining the range to constrain initial points to if *trim.init.range* is *TRUE*.

**1.7.2 Generating the remaining initial points:**

More points are then added until the correct number of points for each species is reached. The method used to generate a new point is controlled with *new.pt.meth.stg.2*, and can either be randomly sampled from the input raster, or placed *D* distance away from an existing conspecific point, where *D* is either sampled with replacement from observed within-species distances or sampled from the empirical distribution function of observed distances (the default). As above, after each point is added, distances are checked to ensure they are within the observed range of distances. If they are not, they are discarded and a new iteration is started. As with *iter.max.stg1*, *iter.max.stg2* sets an upper limit on the number of iterations without improvement, if this is reached the algorithm restarts at stage 1.

*iter.max.stg2*

The maximum number of iterations with no improvement allowed when generating the initial full set of points.

*new.pt.meth.stg.2*

The method for generating the initial full set of points. Either *"sample"*, which samples a random point from the raster; *"dist.obs"* which randomly chooses an existing point and places a new point of the same species *D* distance away, where *D* is sampled from the observed within-species distances for that species; or *"dist.dens"* (recommended) which is similar to dist.obs but *D* is sampled from a density object constructed from the empirical distribution function of observed within-species distances for that species.

**1.7.3 Iteratively improving initial points:**

In the iterative improvement procedure, the number of occurrence points to be replaced per iteration can be set with *switch.n* (one is optimal for our test datasets, but higher values could be more efficient for other datasets). The method of generating new points can be changed as above with *new.pt.meth.stg.3*. The discrete version of the KL divergence is used, in which values are divided into bins, the number of bins can be set using the *break.num* option, which may have a minor effect on performance in some cases.

*new.pt.meth.stg.3*

As with *new.pt.meth.stg.2* but for the iterative improvement stage.

*switch.n*

The number of changes which are made in each iteration.

*break.num*

Integer specifying the number of breaks (to delineate bins) for the KL calculation. Defaults to 20, but this is arbitrary.

**Appendix 2: Dataset preparation and running *fauxcurrence*.**

We tested the method on seven species occurrence datasets from Sulawesi, Indonesia, each including between one and seven species of a single genus. Sulawesi was chosen because a diverse collection of test datasets were available, and its unusual geography makes one of the current leading methods for testing whether niche overlap differs from null expectations, the rotation and translocation method (Nunes & Pearson, 2017), unsuitable (see main text). The datasets were phylogenetically diverse, including plants (*Cyrtandra*, Gesneriaceae, Lamiales: 6 species), lizards (*Cyrtodactylus*, Gekkonidae, Squamata: 1 species; *Draco*, Agamidae, Squamata: 3 species; *Sphenomorphus*, Scincidae, Squamata: 2 species), damselflies (*Libellago*, Chlorocyphidae, Odonata: 4 species; *Rhinocypha*, Chlorocyphidae, Odonata: 2 species) and frogs (*Limnonectes*, Dicroglossidae, Anura: 4 species). Furthermore, they included species with diverse range sizes, from *Cyrtodactylus jellesmae*, with a range that covers most of Sulawesi, to *Limnonectes microtympanum* which is restricted to a small region of South Sulawesi. We used a raster of Sulawesi with 2.5 minute resolution (i.e each raster cell was approximately 4.65 km across), and the occurrence datasets were filtered such that there was a maximum of one occurrence per species in each cell and only species with 10 or more observations following this were included. Because of the unusual geography of Sulawesi, which includes several large peninsulas, we used overland distance for our inter-point distances calculations. We ran all null model replicates until there had been no improvement in KL-divergence for 10,000 iterations. *Fauxcurrence* analyses were conducted in R version 3.6.0 on the Supercomputing Wales cluster which runs on a Red Hat Enterprise Linux Server 7.8 operating system. Each replicate ran on a single core (Intel Xeon Gold 6148) but replicates were parallelised using the foreach package. *Fauxcurrence* was also tested on a MacBook Pro laptop (2019; 2.8 GHz Intel Core i7; 16 GB RAM).

**Appendix 3: Species distribution modelling and niche overlap methods**

We used *Maxent* v. 3.4.1 (Phillips, Anderson, & Schapire, 2006) to build Species Distribution Models (SDMs) for all species using the 19 *BIOCLIM* climate variables and altitude from the *WorldClim1* database (Hijmans, Cameron, Parra, Jones, & Jarvis, 2005) at 2.5 minute resolution as predictor variables. We first selected species-specific settings for the observed datasets following following (Galante et al., 2018) using the *ENMevaluate* function in the

*ENMeval R* package v. 0.3.0 (Muscarella et al., 2014) using regularisation multiplier values between 0.5 and 4 with a step size of 0.5, the “checkerboard2” method of data partitioning, 2000 background points from across Sulawesi, and the following feature classes: "L", "LQ", "H", "LQH", "LQHP" and "LQHPT". *Maxent* was then run on the observed data and all null models for each species using the optimised settings. To determine if SDMs for the observed data had a significantly better fit than the null models, we compared area under the receiver operating characteristic curve (AUC) in SDMs built from observed data to those from the 1,000 replicates from each of the applicable null models. We calculated one-tailed *P*-values as the fraction of null SDM AUCs which were greater than or equal to the observed AUC using the *ecdf* function in *R*. For datasets with more than one species, we then calculated two measures of niche overlap, Schoener’s *D* (Schoener, 1968) and Warren’s *I* (Warren, Glor, & Turelli, 2008), between all pairs of congeneric species in the observed and null datasets using the *calc.niche.overlap* function in the *ENMeval R* package. Because the *Intra* null model ignores between-species distances, it has the potential to simulate occurrence points with far greater range overlap than is found in the observed data. For this reason, it is not appropriate for testing niche overlap, and we only calculated niche overlap for *Inter* and *Inter-sep* models. To test for significant differences between observed and null *I* and *D*-statistics for each species pair, we calculated two-tailed *P*-values as the fraction of null statistics which were closer to the $0.5^{th}$ quantile of all null statistics than the observed value, again using the *ecdf* function. For both tests we considered *P*-values $< 0.05$ to be significant.

### Supplementary figures

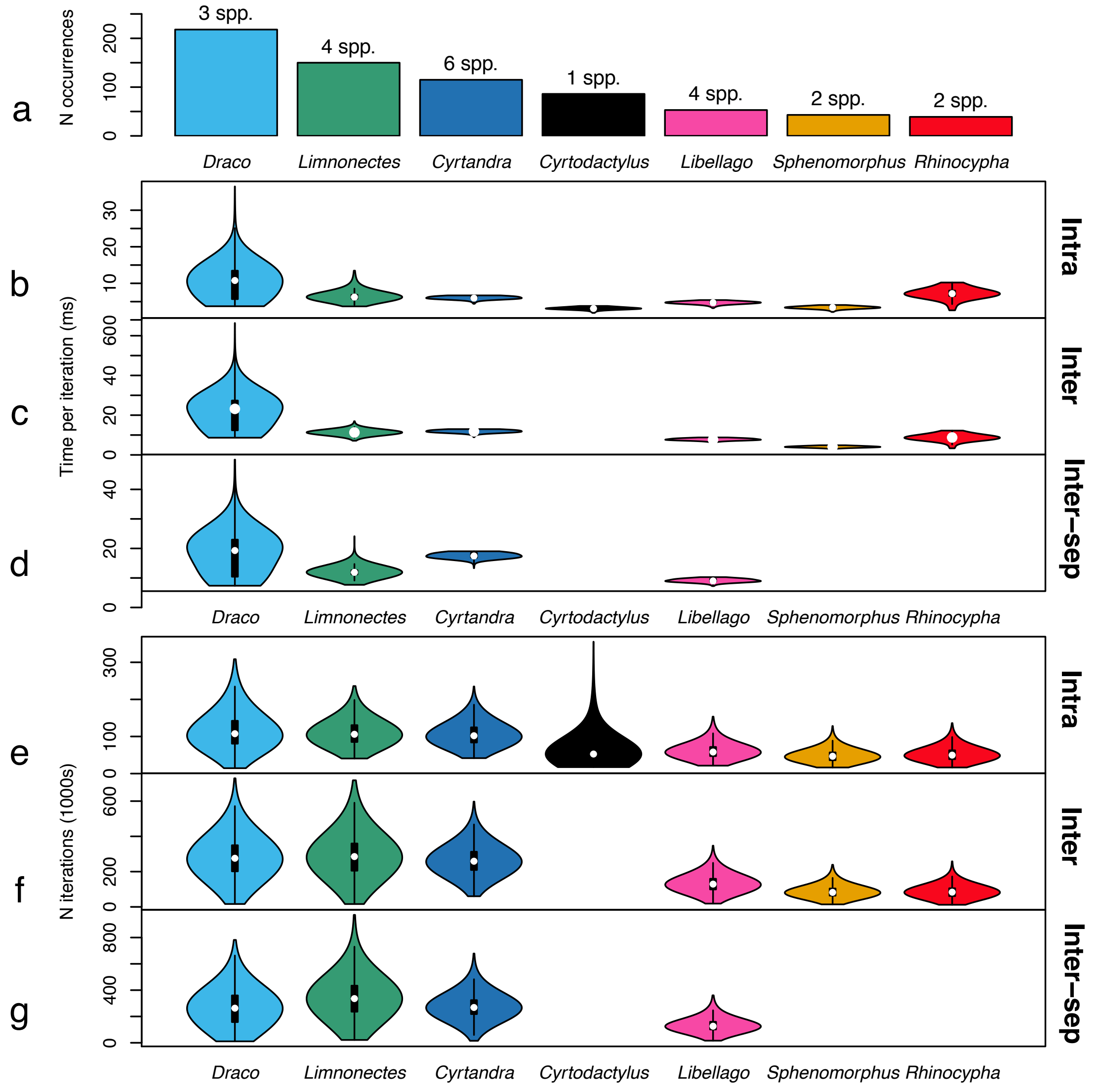


Figure S1. Computational efficiency of the null models. Bar plots show number of occurrences per dataset, with number of species shown above each bar (a). Violin plots: mean computation time per iteration (b-d) and number of iterations (e-g) across the 1000 replicates of each dataset/model combination.

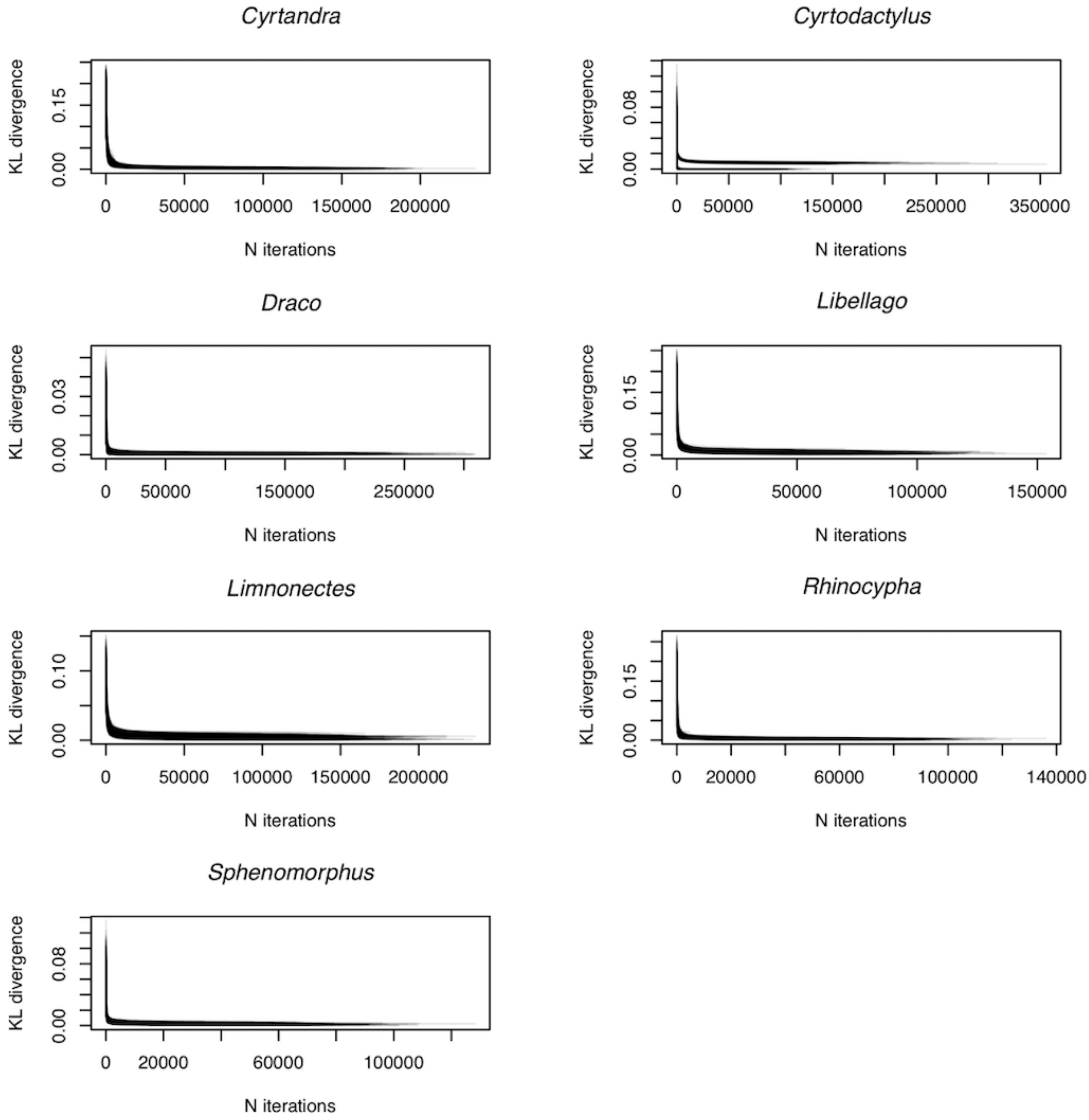


Figure S2. Minimisation of Kullback-Leibler (KL) divergence across iterations using the *Intra* model. For each dataset, the reduction in KL as the algorithm progresses for all 1,000

replicates is shown as overlaid, semi-transparent lines.

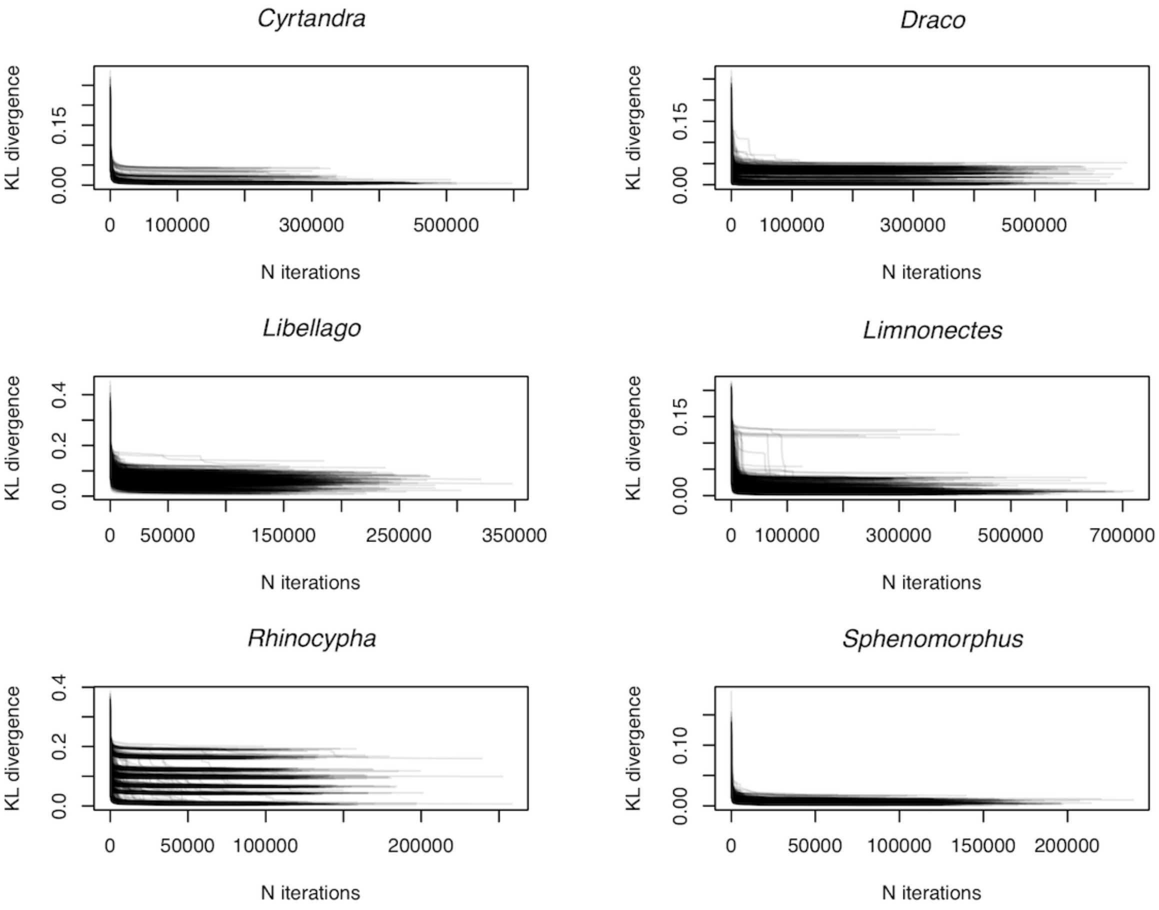


Figure S3. Minimisation of Kullback-Leibler (KL) divergence across iterations using the *Inter* model. Plotted as in Fig. S1.

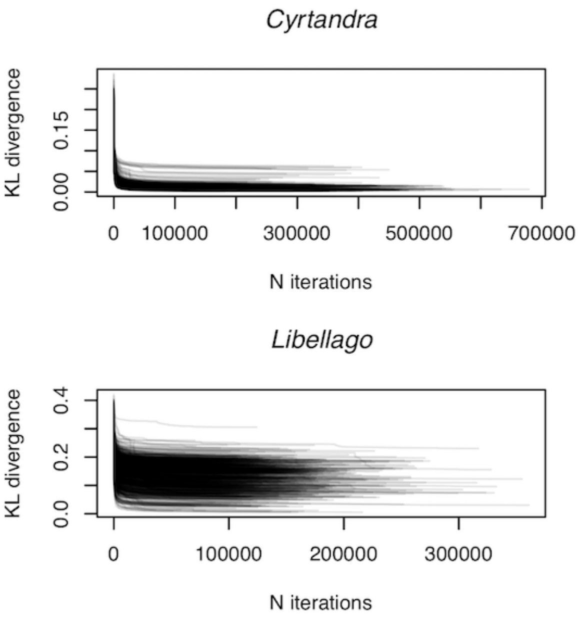


Figure S4. Minimisation of Kullback-Leibler (KL) divergence across iterations using the *Inter-sep* model. Plotted as in Figs. S1-2.

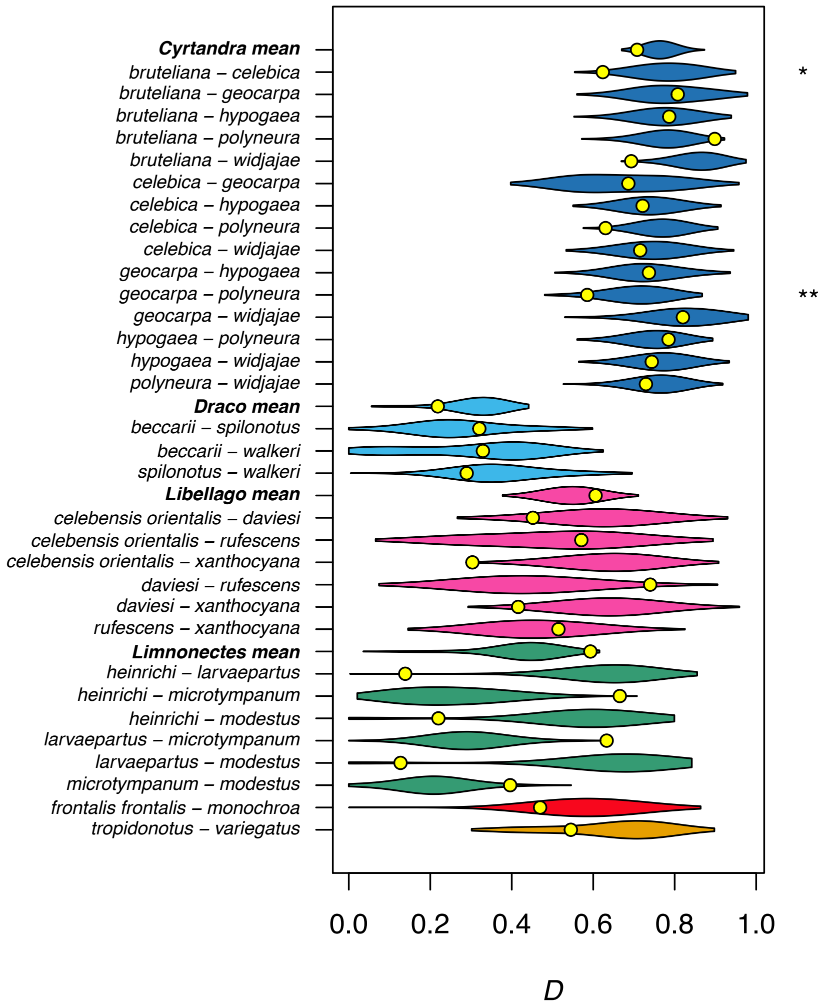


Figure S5. Comparison between observed and null niche overlap using the "*Inter*" null model. Violin plots show the density of Schoener's $D$ across null models for each pair of congeneric species and the mean for genera with more than two comparisons. Observed values are shown by yellow points. Violins are coloured by genus as in Figs. 2-3 (main text), species names are to the left of the plot, and stars to the right of the figure show statistical significance ("*" = $P < 0.05$; "**" = $P < 0.01$).

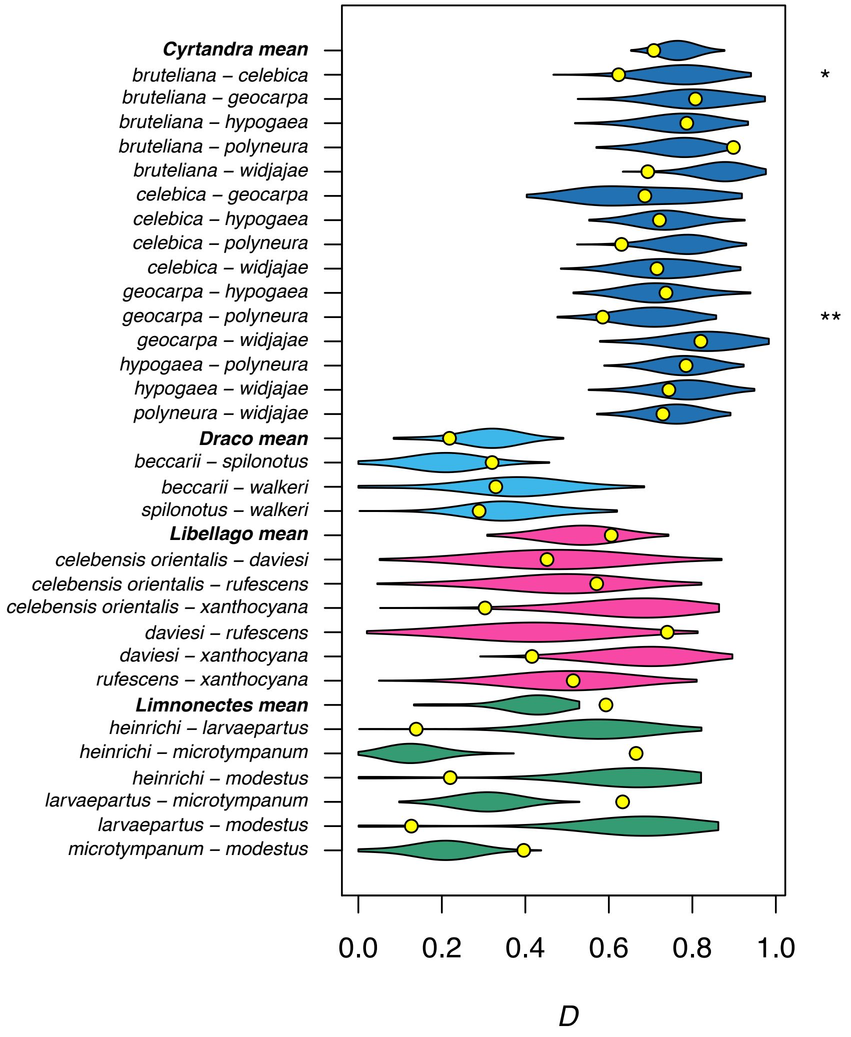


Figure S6 Comparison between observed and null niche overlap using the "*Inter-sep*" null model. Violin plots show the density of Schoener's *D* across null models for each pair of congeneric species and the mean for each genus with more than two comparisons. Observed values are shown by yellow points. Violins are coloured by genus as in Figs. 2-3 (main text), species names are to the left of the plot, and stars to the right of the figure show statistical significance ("*" = $P < 0.05$; "**" = $P < 0.01$).

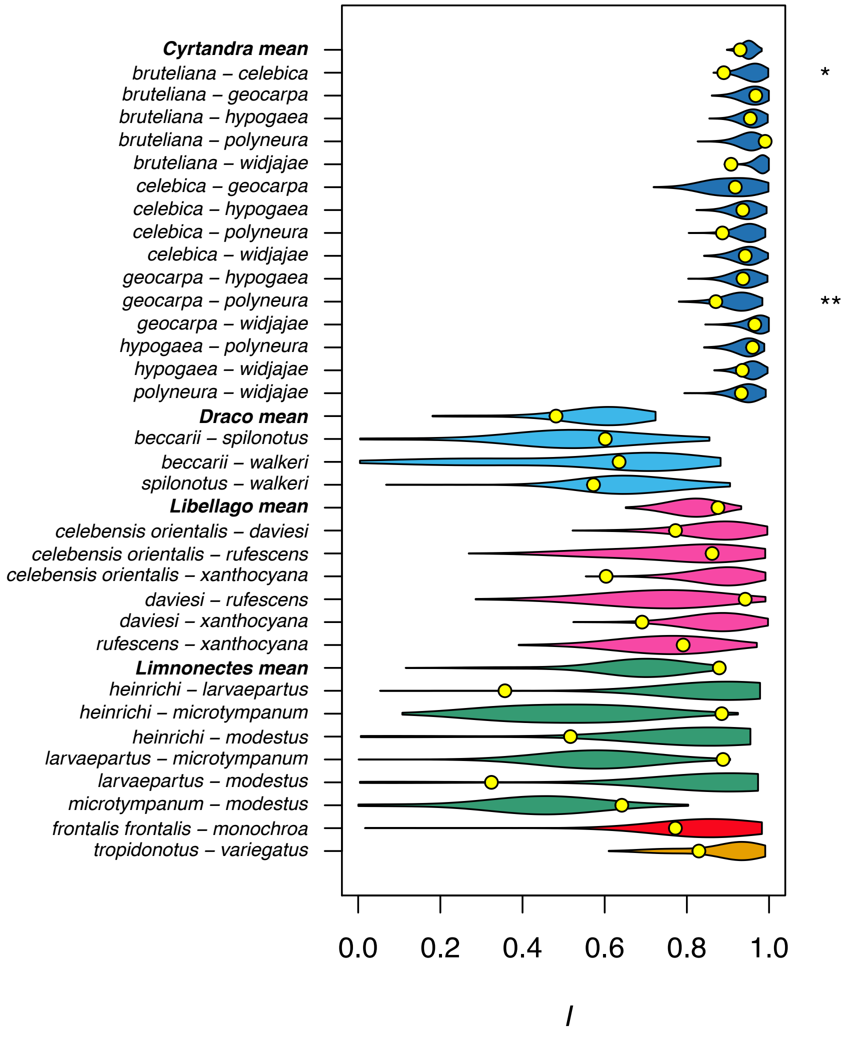


Figure S7. Comparison between observed and null niche overlap using the "*Inter*" null model. Violin plots show the density of Warren's *I* across null models for each pair of congeneric species and the mean for each genus with more than two comparisons. Observed values are shown by yellow points. Violins are coloured by genus as in Figs. 2-3 (main text), species names are to the left of the plot, and stars to the right of the figure show statistical significance ("*" = $P < 0.05$; "**" = $P < 0.01$).

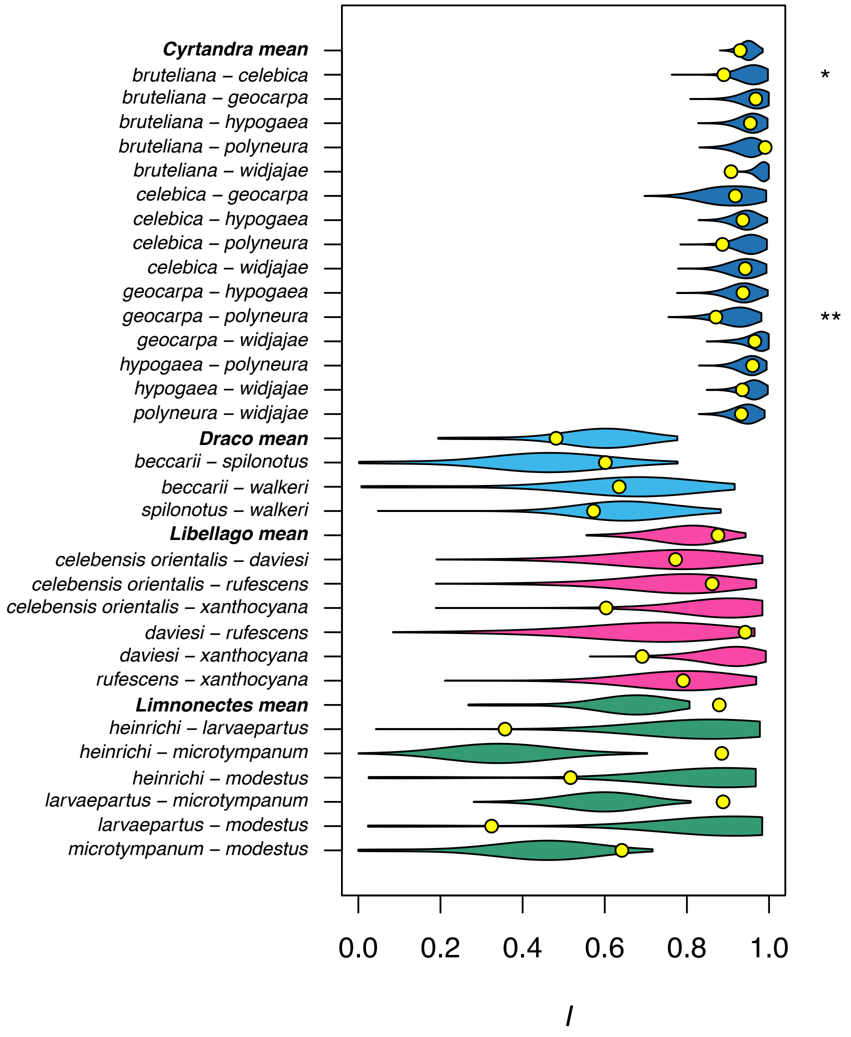


Figure S8. Comparison between observed and null niche overlap using the "*Inter-sep*" null model. Violin plots show the density of Warren's *I* across null models for each pair of congeneric species and the mean for each genus with more than two comparisons. Observed values are shown by yellow points. Violins are coloured by genus as in Figs. 2-3 (main text), species names are to the left of the plot, and stars to the right of the figure show statistical significance ("*" = $P < 0.05$; "**" = $P < 0.01$).

**Supplementary tables**

Table S1. Results of multiple regression analysis to determine if number of occurrences and number of species significantly predicted mean time per iteration or mean number of iterations in the *Intra* and *Inter* models. There were too few datasets used with the *Inter-sep* model for multiple regression analysis, so it isn't included. Abbreviations are as follows: $R^2$(adj) = adjusted $R^2$ for the model; $P$ = $P$-value; $F$ = $F$-statistic for individual predictors; N.occ = number of occurrences; N.spp = number of species; Int = the interaction between number of occurrences and number of species; time = mean time per iteration; N iter = mean number of iterations.

| Null model | Dependent variable | $R^2$ (adj) | $P$ | $F$ N.occ | $P$ N.occ | $F$ N.spp | $P$ N.spp | $F$ Int | $P$ Int |
|---|---|---|---|---|---|---|---|---|---|
| *Intra* | time | 0.038 | 0.476 | 3.215 | 0.171 | 0.019 | 0.899 | 0.006 | 0.945 |
| *Inter* | time | 0.670 | 0.191 | 12.848 | 0.070 | 0.059 | 0.831 | 0.255 | 0.664 |
| *Intra* | N iter | 0.978 | 0.002 | 246.559 | <0.001 | 41.699 | 0.008 | 5.031 | 0.111 |
| *Inter* | N iter | 0.934 | 0.039 | 60.564 | 0.016 | 11.773 | 0.075 | 1.332 | 0.368 |
